# The Xer activation factor of TLCΦ expands the possibilities for Xer recombination

**DOI:** 10.1101/2021.09.09.459576

**Authors:** Solange Miele, Justine Vergne, Christophe Possoz, Françoise Ochsenbein, François-Xavier Barre

**Affiliations:** Institute for Integrative Biology of the Cell (I2BC), Université Paris-Saclay, CEA, CNRS, Université Paris Sud, 1 Avenue de la Terrasse, 91198 Gif-sur-Yvette, France

## Abstract

Many mobile elements take advantage of the highly-conserved chromosome dimer resolution system of bacteria, Xer. They participate in the transmission of antibiotic resistance and pathogenicity determinants. In particular, the toxin-linked cryptic satellite phage (TLCΦ) plays an essential role in the continuous emergence of new toxigenic clones of the *Vibrio cholerae* strain at the origin of the ongoing 7^th^ cholera pandemic. The Xer machinery is composed of two chromosomally-encoded tyrosine recombinases, XerC and XerD. They resolve chromosome dimers by adding a crossover between sister copies of a specific 28 base pair site of bacterial chromosomes, *dif*. The activity of XerD depends on a direct contact with a cell division protein, FtsK, which spatially and temporally constrains the process. TLCΦ encodes for a XerD-activation factor (XafT), which drives the integration of the phage into the *dif* site of the primary chromosome of *V. cholerae* independently of FtsK. However, XerD does not bind to the attachment site (*attP*) of TLCΦ, which raised questions on the integration process. Here, we compared the integration efficiency of thousands of synthetic mini-TLCΦ plasmids harbouring different *attP* sites and assessed their stability *in vivo*. In addition, we compared the efficiency with which XafT and the XerD activation domain of FtsK drive recombination reactions *in vitro*. Taken together, our results suggest that XafT promotes the formation of synaptic complexes between canonical Xer recombination sites and imperfect sites.

## INTRODUCTION

Repair by homologous recombination can lead to the formation of chromosome dimers when the chromosomes are circular, as is generally the case in bacteria and archaea. Chromosome dimers physically impede the segregation of genetic information. They are resolved by the addition of a crossover at a specific locus, *dif*, by a highly conserved <u>c</u>hromosomally <u>e</u>ncoded tyrosine <u>r</u>ecombination (Xer) machinery (1).

Many mobile elements take advantage of the high conservation of the Xer machinery (1). Indeed, it was initially characterized as a multicopy plasmid dimer resolvase (2, 3). Diverse <u>I</u>ntegrative <u>M</u>obile <u>E</u>lements exploiting <u>X</u>er (IMEX) were subsequently discovered, including phages and genetic islands that harbour a *dif*-like attachment site (*attP*) and integrate into the *dif* site of one of the chromosomes of their host (4, 5). Plasmids and IMEX participate in the acquisition of antibiotic resistance and pathogenicity determinants. In particular, the principal virulence factor of *Vibrio cholerae*, <u>c</u>holera <u>t</u>o<u>x</u>in, is encoded in the genome of a lysogenic <u>ph</u>age (CTXΦ), which exploits Xer for integration (6, 7). Several other IMEX contribute to the continuous emergence of new toxigenic clones of the *V. cholerae* strain at the origin of the ongoing 7^th^ cholera pandemic (8–11). Foremost among those is a <u>t</u>oxin-<u>l</u>inked <u>c</u>ryptic satellite <u>ph</u>age (TLCΦ) whose integration is thought to correct the *dif* site of the primary chromosome of non-toxigenic environmental *V. cholerae* strains, *difA*, into a site suitable for the integration of CTXΦ, *dif1* (Figure 1A, (10, 11)).

**Figure 1.**
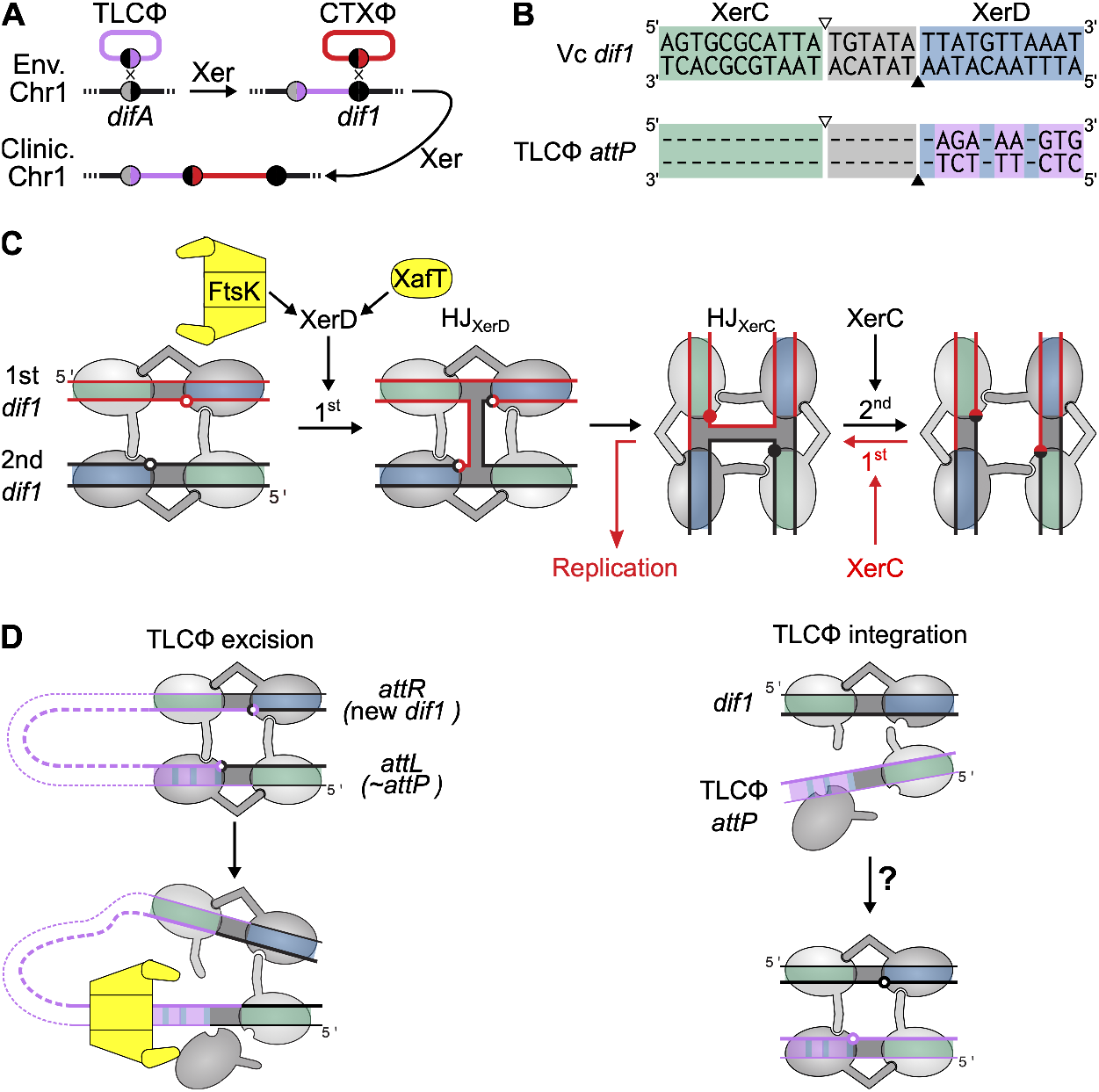
Xer recombination. (A) Toxigenic conversion of *V. cholerae*. Env. and Clinic. Chr1: primary chromosome of environmental and clinical *V. cholerae*. (B) TLCΦ *attP* and *V. cholerae dif1* sequence. Grey: central region; Green: XerC arm; Blue: XerD arm; Pink: TLCΦ non-canonical bp; white and black triangles: XerC and XerD cleavage sites. (C) Xer recombination pathways. Black arrows: conventional recombination pathway; Red arrows: non-conventional asymmetric recombination pathway. The XerD and XerC cleavage points are depicted by empty and filled disks. (D) TLCΦ excision/integration balance. Left: FtsK dismantles the excision complex when it translocates on the prophage DNA; Right: the non-canonical XerD-arm of TLCΦ *attP* abolishes binding of XerD.

In *V. cholerae*, as in most bacteria, the Xer machinery is composed of two closely related recombinases, XerC and XerD (Figure S1A). The *dif* sites are composed of two partially-palindromic 11 base pair (bp) XerC and XerD binding arms separated by a short 6 bp central region (Figure 1B and S1B). XerC and XerD each catalyse the cleavage and transfer of a specific pair of DNA strands, which are referred to as the top and bottom strands, respectively (Figure 1B). Chromosome dimer resolution follows the conventional recombination pathway of tyrosine recombinases, with the cleavage and transfer of one pair of strands leading to the formation of an obligatory Holliday junction (HJ) intermediate that is subsequently resolved into crossover by the exchange of a second pair of strands (Figure 1C). The first pair of strands is catalysed by XerD and the second by XerC (Figure 1C, (12, 13)). The process is under the control of a DNA translocase anchored in the cell division septum, FtsK, which activates XerD by a direct contact with its C-terminal domain, FtsKγ (Figure 1C, (12, 14–16)). Hence, Xer recombination is normally restricted to the time of cell division and to synaptic complexes located in the proximity of the cell division apparatus (17–20). However, many IMEX, including CTXΦ, integrate via a non-conventional FtsK-independent recombination pathway: they exploit the low basal ability of XerC to catalyse the formation of HJs, which are subsequently resolved by replication (Figure 1C, (8, 9, 21)). Dimers of the ColE1 multicopy plasmid are resolved by a similar process (22). In contrast, TLCΦ integration follows the same recombination pathway as chromosome dimer resolution (10). However, the process escapes the FtsK control (10) because <u>T</u>LCΦ encodes for its own <u>X</u>erD <u>a</u>ctivation <u>f</u>actor, XafT (Figure 1C, (23)).

Eight base pairs of the XerD-arm of TLCΦ *attP* deviate from the canonical XerD-arm of *dif1* and *difA* (Figure 1B and S1B). It was proposed to prevent undesired FtsK-driven prophage excision, at least in part because FtsK can dismantle non-canonical synaptic complexes when it translocates on the genome of the integrated IMEX (Figure 1D, (24)). However, the XerD arm of TLCΦ *attP* abolishes XerD binding, which questioned the possibility for integration (Figure 1D, (10, 11, 23)).

Here, we analysed the influence of the sequence of the XerD arm of TLCΦ *attP* on FtsK- and XafT-driven recombination reactions by comparing the integration efficiency of thousands of synthetic mini-TLCΦ plasmids harbouring *attP* sites with differing XerD-arms and assessing their stability *in vivo*. In addition, we analysed the efficiency with which XafT and FtsKγ drive recombination reactions between two *dif1* sites and between *dif1* and TLCΦ *attP*. Taken together, our results suggest that XafT promotes the formation of synaptic complexes between *dif* sites and imperfect Xer recombination sites.

## RESULTS

### Methodology for parallel monitoring of the integration efficiency and stability

We developed a methodology based on Next Generation Sequencing (NGS) to analyse the impact of the sequence of the XerD-arm of TLCΦ *attP* on the integration efficiency and stability of the phage. In brief, we generated three pools of 65,536 (4^8^) degenerate *attP* sites using synthetic oligonucleotides carrying 8 degenerate bases at different positions of the XerD-arm of TLCΦ *attP* as templates (Figure 2A). Two of the pools, referred to as n_8_gtg and tagn_8_ on the basis of their top strand, were designed to explore the influence of the eight innermost and outermost positions of the XerD-arm. Results were completed with a pool harbouring degenerate bases in both the inner and outer part of the XerD-arm, n_5_a_2_n_3_g. The pools were cloned in place of the *attP* site of previously designed XafT^+^ and XafT^-^ conjugative suicide mini-TLCΦ plasmids (23). NGS analysis showed that each resulting mini-TLCΦ library contained over 99.5% of the XerD-arm sequence motifs covered by the pool from which it originated, with a median copy number in the order of 15 per million reads (Figure 2A). It further showed that the XafT^+^ and XafT^-^ mini-TLCΦ libraries contained over 99.9% and 99.8% of the 190,528 XerD-arm sequence motifs covered by the three combined degenerate pools, respectively. The mini-TLCΦ libraries were conjugated in two N16961 reporter strains harbouring an *E. coli lacZα-dif1-lacZβ* gene fusion at the natural integration locus of TLCΦ or at the *lacZ* locus (Figure S2, (10)). As the central region and XerC-binding arm of TLCΦ *attP* and *dif1* are identical, the *attR* and *attL* site resulting from the integration of the mini-TLCΦ plasmids are identical to their *attP* site and *dif1*, respectively. Thus, the integration and excision frequencies of the mini-TLCΦ plasmids reflect the efficiency of intermolecular and intramolecular recombination reactions between the same two sites, respectively. The global frequency of *dif1*-integration events of each mini-TLCΦ library, *f*, was measured with a blue/white screen (Figure 2B). The relative proportion of each *attP* sequence motif in the genomic DNA of the recipient cells*, P_int_*, was determined by NGS using a primer binding in *E. coli lacZ* and a primer binding in TLCΦ (Figure 2B). The integration frequency of the corresponding mini-TLCΦ plasmids was then estimated as *f_int_* = *f* × *P_int_*. The reporter strains were further engineered to place production of XerC and XerD under the control of the arabinose promoter in order to prevent Xer-mediated excision after integration (Figure 2B, (25)). The stability of the integrated plasmids was estimated by analysing the proportion of each *attP* sequence motif in the cell population after growth in the presence of arabinose, *P_exc_* (Figure 2B).

**Figure 2.**
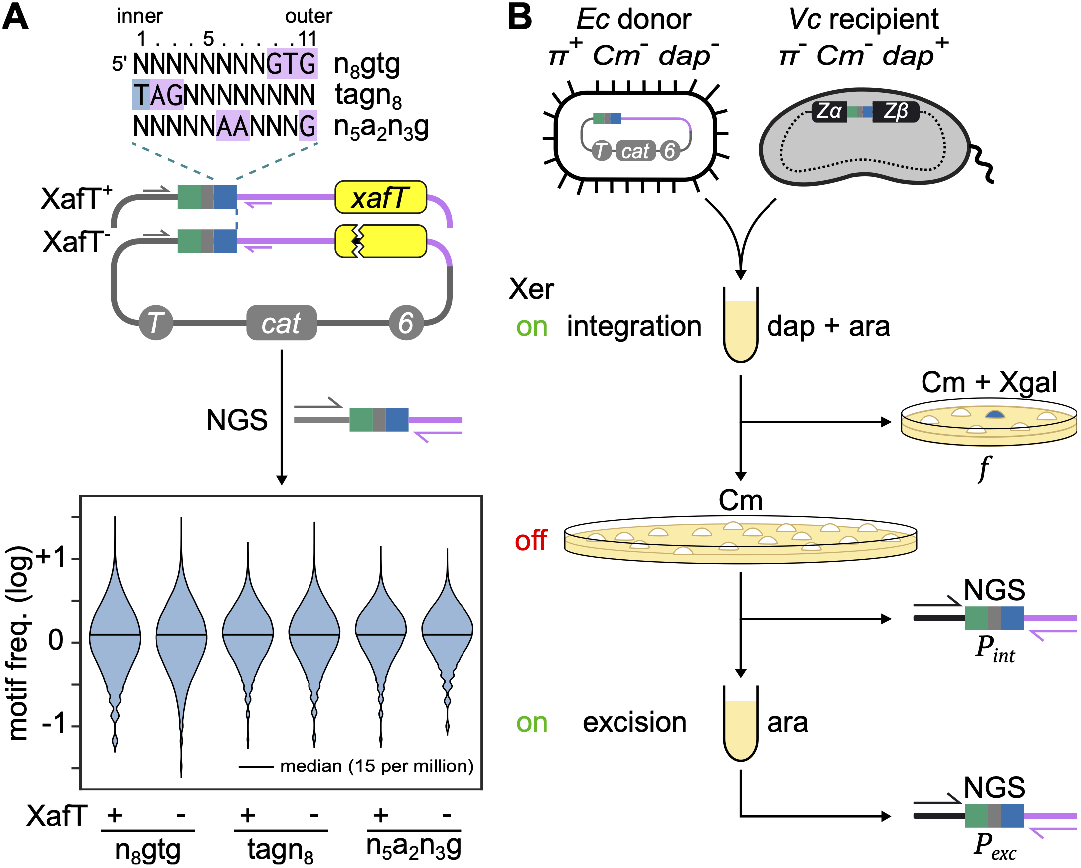
Parallel monitoring of integration and stability. **(A)** Mini-TLCΦ plasmid libraries. Top panel: top strand of the degenerate *attP* motifs. Legend as in Figure 1. Middle panel: scheme of XafT^+^ and XafT^-^ mini-TLCΦ plasmids. Pink: TLCΦ DNA; Grey: plasmid DNA. Yellow rectangle: *xafT* gene; sawed lines: stop mutation; T: RP4 transfer origin; 6: *pir*-dependent replication origin; *cat*: chloramphenicol resistance gene. Plasmid-specific P5 and TLCΦ-specific P7 adaptor primers used for next generation sequencing (NGS) are indicated by grey and pink arrows, respectively. Bottom panel: distribution frequency of the motifs. **(B)** Integration and excision assays. On and off: growth in the presence or absence of arabinose. π: R6K replication initiator; dap: diaminopimelic acid; Cm: chloramphenicol; Xgal: X-gal; *Zα and Z*β: *E. coli lacZ* gene. The *lacZα*-specific P5 adaptor primer is indicated by a black arrow.

### FtsK-driven integration is restricted to the sites that most resemble *dif1*

To explore the influence of the sequence of the XerD-arm of TLCΦ *attP* on the efficiency of FtsK-driven integration events, we conjugated the XafT^-^ mini-TLCΦ libraries in the *V. cholerae* recipient strain harbouring *dif1* at its natural locus (Figure 3, FtsK panel). The global integration frequency of the XafT^-^ n_8_gtg, tagn_8_ and n_5_a_2_n_2_g mini-TLCΦ libraries was 1000-fold lower than that of a XafT^-^ mini-TLCΦ plasmid harbouring *dif1* (Figure 3A, FtsK panel). NGS further revealed that 2.5% of the different possible *attP* sites comprised in the three XafT^-^ libraries could be integrated (Figure 3A, FtsK panel).

**Figure 3.**
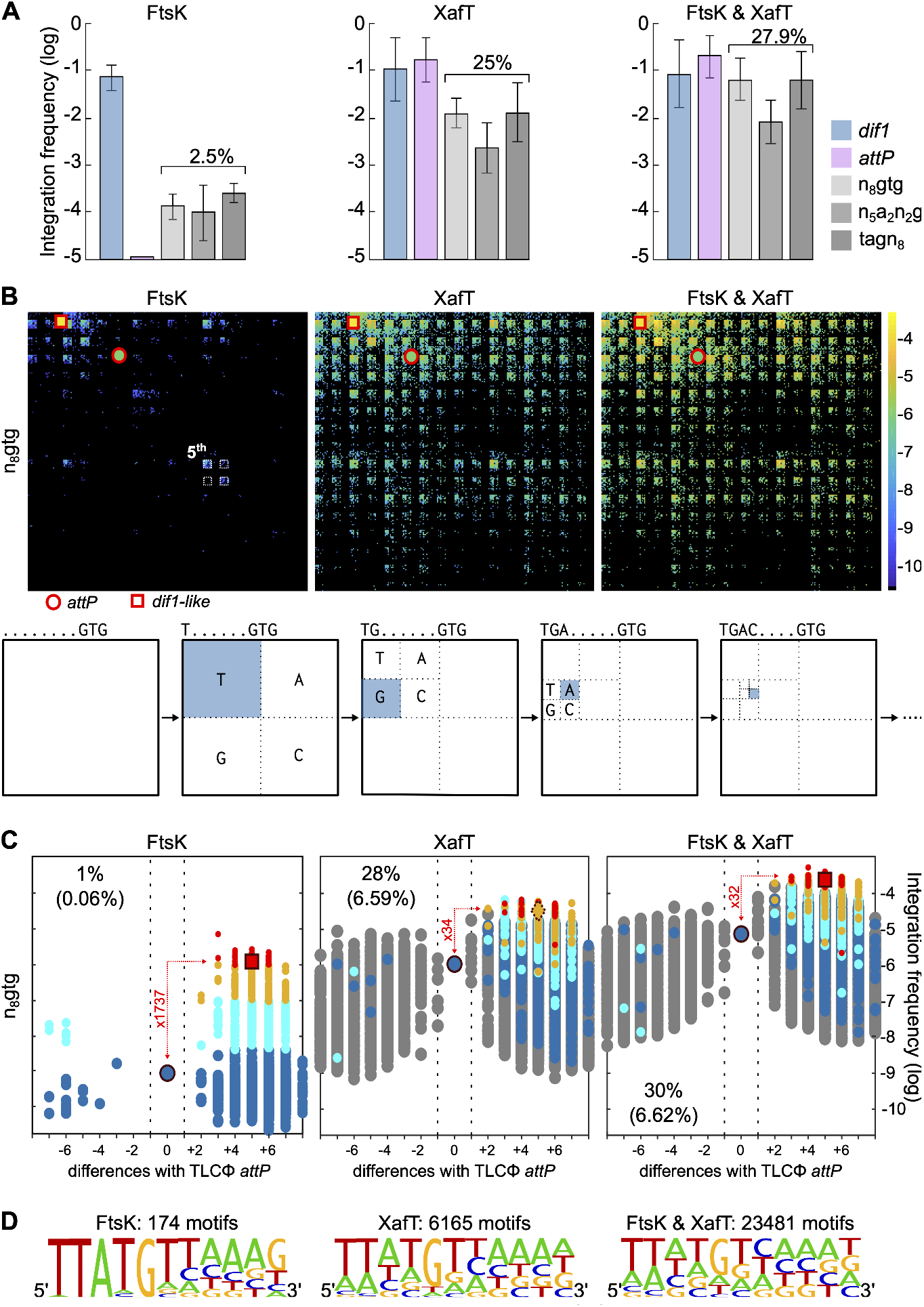
Suicide mini-TLCΦ plasmid integration. **(A)** Integration frequencies of mini-TLCΦ plasmids harbouring *dif1* or TLCΦ *attP*, and global integration frequencies (*f*) of n_8_gtg, n_5_a_2_n_2_g and tagn_8_ plasmid libraries. FtsK panel: XafT^-^ plasmids conjugated in a strain harbouring *dif1* at its natural locus; XafT panel: XafT^+^ plasmids conjugated in a strain harbouring *dif1* at the *lacZ* locus; FtsK & XafT panel: XafT^+^ plasmids conjugated in a strain harbouring *dif1* its natural locus. Mean and standard deviations of at least 3 independent assays. **(B)** 2D maps showing the relative integration frequency of the different possible n_8_gtg sequences (*f_int_*). The scheme below the 2D maps indicates how specific horizontal and vertical coordinates are assigned to each degenerate motif. The positions of TLCΦ *attP* and of the sequences most similar to *dif1* are highlighted. **(C)** Butterfly plots showing the relative integration efficiency of the different possible n_8_gtg sequences. X-axis: number of changes from TLCΦ *attP* (from 0 to 8). + and - values indicate whether the changes render the site more similar to *dif1* or not. The proportion of sequences falling in the - group is indicated. Their contribution to the global frequency of integration of the library is shown between brackets. A red, orange, cyan and blue colour code was assigned to the *attP* sites from the FtsK panel whose integration frequencies was higher than 10^-6^, 10^-7^, 5 10^-8^ or lower than 5 10^-8^, respectively. The corresponding sites in the XafT and Ftsk & XafT panels were highlighted with the same colour code, with sites absent in the FtsK panel shown in grey. The difference between the mean frequency of integration of red motifs and the integration frequency of TLCΦ *attP* is indicated in red**. (D)** Number of different n_8_gtg, n_5_a_2_n_2_g and tagn_8_ sequences with an at frequencies higher than 10^-6^. The sequence logo shows the frequency of each base at the degenerate positions.

To visualise the sequence bias of the XerD arm of the *attP* sites of the integrated plasmids, we attributed unique x and y coordinates to each XerD arm motif based on its sequence (Figure 3B, bottom scheme). We then drew two-dimensional maps (2D-maps) by colouring each x and y position based on the integration frequency of the corresponding *attP* site, from dark blue to bright yellow. The position of non-recovered *attP* sites were coloured in black. The density of the coloured positions highlighted the limited number of XerD-arm sequences that could be integrated by FtsK (Figure 3B and S3A, FtsK panels). A single small yellow tile was visible on the n_8_gtg and n_5_a_2_n_2_g 2D-maps, at the coordinates of the 64 <u>TTATGN</u>NNNGTG and 64 <u>TTATG</u>AANNNG top strand sequences, respectively. A larger blue tile was visible on the tagn_8_ 2D-map, which corresponded to the 1024 TAG<u>TGT</u>NNNNN top strand sequence coordinates. The lower frequency of integration and higher number of motifs recovered with the tagn_8_ library showed that inner XerD positions played a more important role on FtsK-driven integration than outer positions.

Results were further analysed using *butterfly* plots, in which y-axis positions indicate the integration frequency of the sites and x-axis positions indicate their number of differences from TLCΦ *attP*, with positive and negative values attributed to sites that are closer from *dif1* than TLCΦ *attP* (Figure 3C and S3B, FtsK panels). While only a third of the *attP* sites of the n_8_gtg, n_5_a_2_n_2_g and tagn_8_ libraries were closer from *dif1* than TLCΦ *attP* (21 070/ 65 536), over 98% of the sites whose integration could be driven by FtsK belonged to + wing of the plots (Figure 3C and S3B, FtsK panel). In addition, those sites accounted for over 99.6% of the integration events (Figure 3C and S3B, FtsK panel). We assigned a red, orange, cyan and blue colour code to the *attP* sites that integrated at *dif1* by an FtsK-driven reaction at frequencies higher than 10^-6^, 10^-7^, 5 10^-8^ or lower than 5 10^-8^, respectively. The red sites comprised 53 n_8_gtg, 61 n_5_a_2_n_2_g and 61 tagn_8_ sequences, which represented a total of 174 different sequences. We calculated the frequency of the four DNA bases at each position of the top strand of the XerD arm of the red sites by pooling together the sequences coming from libraries in which the position was degenerated. In all but the 11^th^ outermost position of the XerD arm, the *dif1* base was the most frequent (Figure 3D, FtsK panel).

Together, those results show that FtsK-driven integration is restricted to the sites that most resemble *dif1*. Nevertheless, the NGS-based integration assay was sensitive enough to monitor the integration frequency of the XafT^-^ mini-TLCΦ plasmid harbouring TLCΦ *attP*, in contrast to the classical blue/white assay (Figure 3, FtsK panels). The integration frequency of TLCΦ *attP* was over a 1000-fold lower than the mean integration frequency of the n_8_gtg and n_5_a_2_n_2_g red *attP* sites, and over a 100-fold lower than the mean integration frequency of the red tagn_8_ *attP* sites (Figure 3C and S3C, FtsK panel).

### XafT drives the integration of any *attP* site harbouring the same 5^th^ XerD-arm bp than *dif1*

Previous works showed that the choreography of chromosome segregation restricts the activity of FtsK to the terminus region of the two *V. cholerae* chromosomes, suggesting that FtsK could not drive integration at the *lacZ* locus (Figure S2, (25–28)). Correspondingly, no *dif1*-specific integration events were observed at the *lacZ* locus with XafT^-^ mini-TLCΦ libraries. Thus, we could explore the influence of the sequence of the XerD-arm of TLCΦ *attP* on the efficiency of XafT-driven integration events by conjugating the XafT^+^ mini-TLCΦ libraries in the *V. cholerae* recipient strain harbouring *dif1* at the *lacZ* locus (Figure 3, XafT panels).

The integration of the XafT^+^ *dif1* and XafT^+^ TLCΦ *attP* plasmids at the *lacZ* locus was as efficient as the integration of the XafT^-^ *dif1* plasmid at the *dif1* locus (Figure 3A, XafT panel). In addition, the global integration frequency of the XafT^+^ n_8_gtg, tagn_8_ and n_5_a_2_n_2_g mini-TLCΦ plasmid libraries was only 10-fold lower than that of the XafT^+^ *dif1* plasmid (Figure 3A, XafT panel). NGS analysis further revealed that over 25% of the different possible *attP* sites comprised in each of the three XafT^+^ mini-TLCΦ libraries could be integrated at the *lacZ* locus (Figure 3A, XafT panel).

The n_8_gtg and n_5_a_2_n_2_g 2D-maps presented a very striking chequered pattern, with the black stripes corresponding to members of the library that lacked a G at the 5^th^ position of the top strand of the XerD-arm (Figure 3B and S3A, XafT panel). Likewise, the checker pattern of the tagn_8_ 2D maps reflected the importance of the nature of the residue at this position. Thus, XafT promoted the integration of 1/4 of the *attP* library, corresponding to those that carried the same residues as *dif1* at the 5^th^ position of the XerD arm.

The colour of the XafT^+^ 2D maps patterns ranged from yellow to green, showing that XafT-driven integration was more efficent than XafT-driven integration (Figure 3B and S3A, XafT panel). In particular, the integration frequency of TLCΦ *attP* jumped from about 10^-9^ to about 10^-6^ (Figure 3C and S3B, XafT panels). In addition, differences in the frequency of *attP* sites from the red and orange categories were alleviated. In particular, the mean frequency of the red n_8_gtg and n_5_a_2_n_2_g *attP* sites was now only 30-fold higher than that of TLCΦ *attP* (Figure 3C and S3B, XafT panel). Furthermore, the integration frequency of many of the newly recovered sites, shown in grey, was higher than 10^-6^. Those sites were not limited to the positive wing of the butterfly plots: XafT drove the integration of about 30% of the n_8_gtg and n_5_a_2_n_2_g mini-TLCΦ and about 20% of the tagn_8_ mini-TLCΦ that belonged to the negative group (Figure 3C and S3B, XafT). As a result, over 6000 different *attP* sites were found integrated at a frequency higher than 10^-6^, with a frequency logo that lost similarity to *dif1* (Figure 3D, XafT panel).

### FtsK does not perturb XafT-mediated integration at the *dif1* locus

XafT ensured the efficient integration of many *attP* sites other than TLCΦ *attP* (Figure 3C and S3, XafT panels). Therefore, we wondered whether TLCΦ *attP* had been selected to avoid any perturbation of FtsK on the integration of the phage at the natural *dif1* locus. To explore this possibility, we analysed the influence of the sequence of the XerD-arm of TLCΦ *attP* on the efficiency of integration of XafT^+^ mini-TLCΦ libraries that were conjugated in the *V. cholerae* recipient strain harbouring *dif1* at its natural locus (Figure 3 and S3, XafT & FtsK panels). The 2D maps and butterfly plots were similar to those obtained when the libraries were conjugated in the *V. cholerae* recipient strain harbouring *dif1* at the *lacZ* locus (Figure 3 and S3, XafT & FtsK panels). The only notable difference was a slightly higher global integration frequency (Figure 3A and S3, XafT & FtsK panel). This is most probaly explained by the growth advantage of the reporter strain carrying *dif1* at its natural locus over the reporter strain carrying *dif1* at it’s the *lacZ* locus, since the later cannot resolve dimers of its primary chromosome (13). As a result, over 20000 different *attP* sites integrated at a frequency higher than 10^-6^, which decreased the similitude between the sequence logo and *dif1* (Figure 3D, XafT & FtsK panel).

### Mini-TLCΦ plasmids with *attP* sites deviating from *dif1* escape FtsK-driven excision

An unusually long IMEX, the gonococcal genomic island (GGI), is integrated at the *dif* site of the chromosome of pathogenic Neisseria species (29, 30). It is flanked by a *dif-like* site with 4 non-canonical bp within the 6 outermost positions of the XerD-arm. The FtsK DNA translocase was shown to strip XerC and XerD bound to this site, thereby preventing GGI excision (24). It suggested the possibility that TLCΦ *attP* was selected to allow XafT-driven integration while avoiding FtsK-driven excision events (Figure 1D). To explore this hypothesis, we monitored the stability of XafT^+^ n_8_gtg, tagn_8_ and n_5_a_2_n_2_g mini-TLCΦ plasmids that were integrated at the *dif1* locus. Production of XerC and XerD did not reduce the number of *attP* sites in the cell population (Figure 4A and S4, butterfly plots). However, there was a significant decrease in the proportion of *attP* sites from the red and orange categories (Figure 4A and S4, bar plots). Most importantly, the frequency of TLCΦ *attP* became higher than the frequency of the *attP* site that most resembled *dif1* (Figure 4A and S4, butterfly plots).

**Figure 4.**
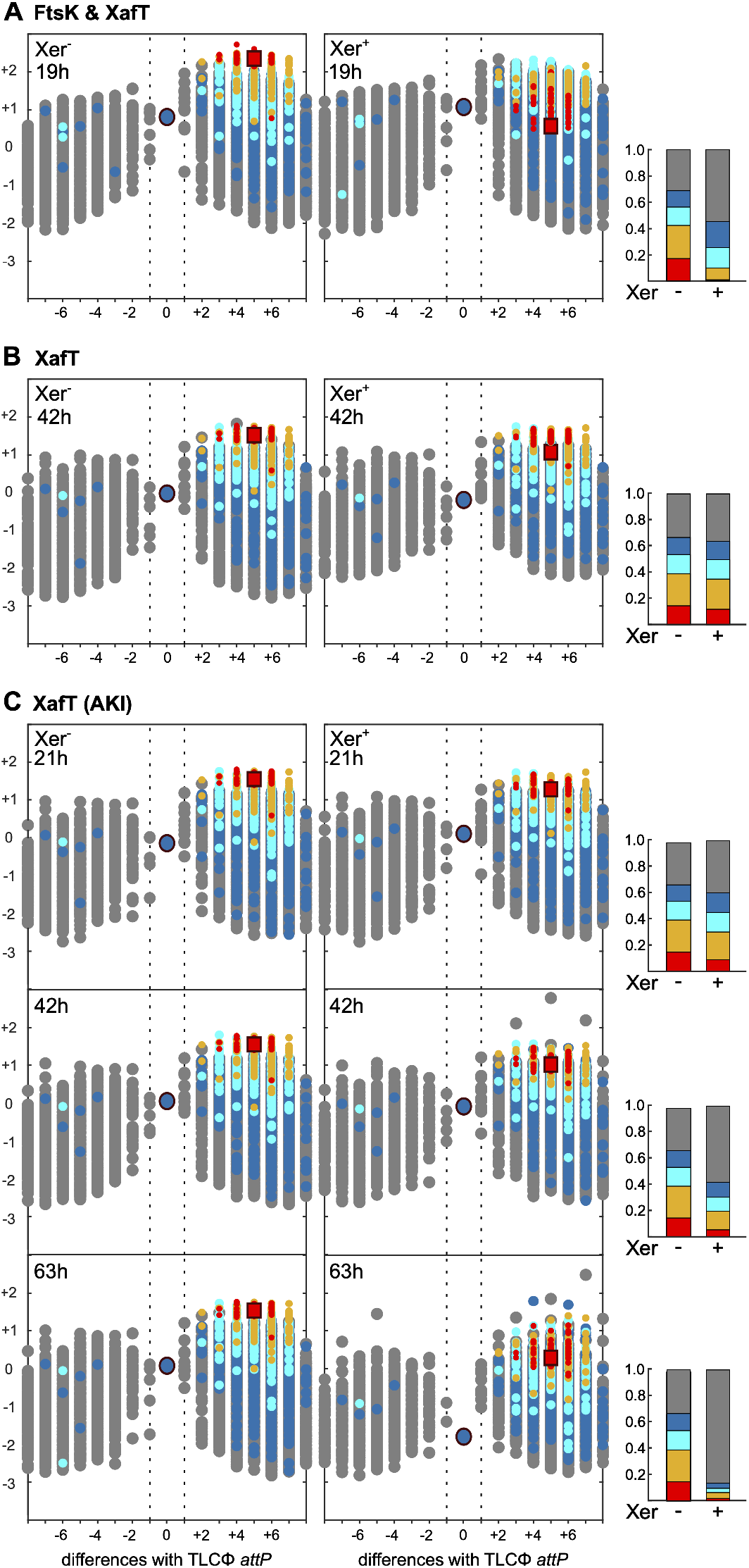
Relative stability of XafT^+^ plasmids harbouring an *attP* site with the n_8_gtg motif. **(A)** Remaining plasmids at the natural *dif1* locus after growth in LB. **(B)** Remaining plasmids integrated at the *lacZ* locus after growth in LB. **(C)** Remaining plasmids integrated at the *lacZ* locus after growth in AKI. Butterfly plots show the frequency of each motif as in Figure 3. Bar plots show the proportion of remaining motifs from the red, orange, cyan, blue and grey categories. Xer^+/-^ indicate whether the inducer (L-arabinose) of XerC and XerD production was added to the growth medium or not.

To determine the relative contribution of FtsK and XafT to the observed excision events, we analysed the stability of XafT^+^ n_8_gtg plasmids integrated at the *lacZ* locus, where FtsK cannot act (Figure S2). After 42h of growth, there were little changes in the frequency of the different *attP* sites (Figure 4B, butterfly plots) and in the relative proportion of the sites from the 5 colour code categories in the cell population (Figure 4B, bar plots). The proportion of the *attP* sites from the red category went down, but the decrease was much less important than the one observed at *dif1* after only 19h of growth. These results suggested that the expression of XafT was repressed after integration under normal laboratory growth conditions. In contrast, the proportion of *attP* sites from the red and orange categories was reduced by half after 42h of growth in AKI, a medium designed to mimic the intestinal environment, which was previously shown to boost the expression of the genes from another *V. cholerae* IMEX, CTXΦ (Figure 4C, bar plots, (31)). Furthermore, *attP* sites from the red, orange, cyan and blue categories represented less than 10% of the total sequence reads after 63h of growth in AKI. In particular, there was a 100-fold decrease in the relative proportion of TLCΦ *attP* (Figure 4C, butterfly plots).

We conclude that FtsK is responsible for the excision of mini-TLCΦ plasmids integrated at the *dif1* locus under normal laboratory growth conditions. However, the excision frequency of most sites remained far lower than the 20% per generation excision frequency of a DNA cassette flanked by two *dif1* sites (25). Thus, many other sites than TLCΦ *attP* could have been selected for stability.

### TLCΦ *att*P/ *dif1* synaptic complexes are rare and/or transient

XafT activates XerD in *trans* via a direct interaction (Figure 5A, (23)). FtsKγ also interacts with XerD (14, 16, 23, 32). However, it acts in *cis* when FtsK is bound on the XerD side of one or the other of the two DNA duplexes engaged in the recombination complex (Figure 5A, (33, 34)). It suggested that FtsK translocation could dismantle TLCΦ *attP/ dif1* synaptic complexes before FtsKγ had had the time to activate XerD, thereby preventing FtsK-driven integration events (Figure 5A, (24)). We reasoned that if this hypothesis was correct, a C-terminal fusion of FtsKγ to XerD, which was previously shown to maximize the efficiency of reactions mediated by the *E. coli* and *N. gonorrhoea* Xer recombinases, should be able to promote the recombination of *dif1* with TLCΦ *att*P *in vitro* (15, 24). As a point of comparison, we reconstituted XafT-mediated reactions using XerC and XerD proteins and an N-terminal fusion of XafT to the maltose binding protein, MBP-XafT (23). We used a short 34-bp synthetic double-stranded DNA (dsDNA) *dif1* fragment and longer dsDNA fragments containing either *dif1* or TLCΦ *attP* as substrates. To facilitate the differentiation of the HJ intermediate, the two crossover products and the substrate, the 5’ and 3’ sides of the bottom strand of the short *dif1* substrate were labelled with cy5 and cy3 fluorescent dyes, respectively (Figure 5B). Incubation of the two *dif1* substrates with XerC and XerD-FtsKγ or with XerC, XerD and MBP-XafT yielded similar amounts of HJs and crossover products, suggesting that FtsKγ promoted recombination as efficiently as XafT when it was fused to XerD (Figure 5C, left panel). Yet, incubation of the short labelled *dif1* substrate and the TLCΦ *attP* substrate with XerC and XerD-FtsKγ yielded barely detectable amounts of HJs and crossover products, invalidating the idea that the low frequency of FtsK-driven integration events could be attributed to the translocase activity of FtsK (Figure 5C, right panel). In contrast, XerC, XerD and MBP-XafT mediated the formation of similarly high amounts of HJs and crossover products between the short labelled *dif1* substrate and the long *dif1* or TLCΦ *attP* substrates (Figure 5B). We conclude that TLCΦ *attP/dif1* synaptic complexes are normally too rare and/or too transient to allow for FtsKγ-driven activation of XerD.

**Figure 5.**
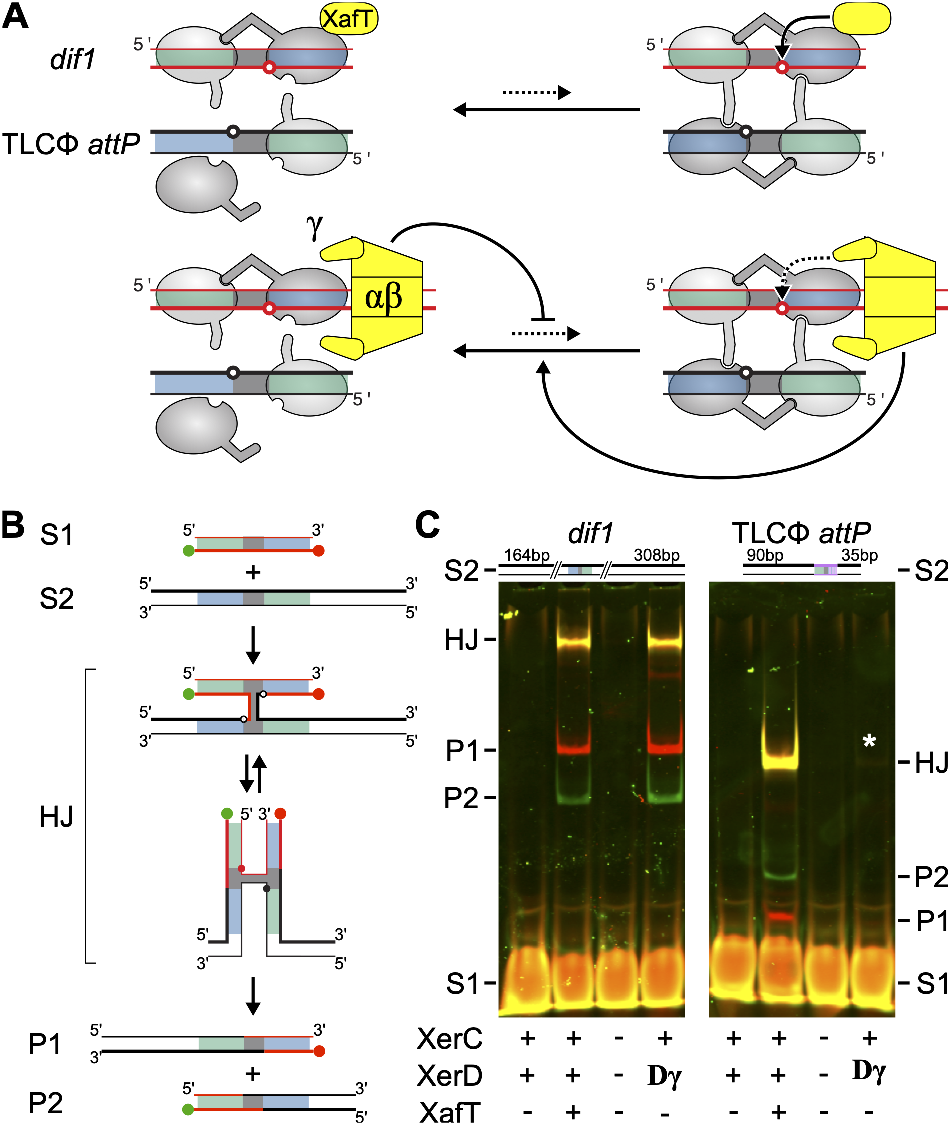
Relative efficiency of XafT- and FtsKγ-driven Xer recombination reactions. **(A)** Scheme of the recombination substrates, the HJ recombination intermediate and the crossover products. Green ball: 3’ Cy3 label; Red ball: 5’ Cy5 label; Grey, Green and Blue rectangles: Central region, XerC-binding and XerD-binding arm, respectively. The DNA strands exchanges by XerC and XerD are depicted as thin and fat lines, respectively. **(B)** XafT- and FtsKγ-promoted recombination reactions. Left panel: recombination of a short labelled *dif1* substrate (S1) and a long non-labelled *dif1* substrate (S2); Right panel: recombination of a short labelled *dif1* substrate (S1) and a long non-labelled TLCΦ *attP* substrate (S2). The S2 substrates are depicted above each gel images. +: *V. cholerae* XerD, *V. cholerae* XerD or MBP-XafT, as indicated; γ: *V. cholerae* XerD-FtsKγ fusion; -: mock buffer of the corresponding purified proteins. A white star indicates the presence of a faint HJ band.

## DISCUSSION

Xer recombination takes place in a nucleoprotein complex consisting of two recombining sites and a pair of each of the two XerC and XerD recombinases (Figure 1). The complex is held together by the affinity of the recombinases to their DNA binding sites and cyclic interactions between their C-terminal domains. However, binding of XerC to *dif* is relatively weaker than binding of XerD (2, 13, 35). Those observations suggested that contacts between XerD and *dif* played a primary role in the assembly and stability of the recombination synapses (2, 13, 35).

In this regard, the efficiency with which TLCΦ integrated in the primary chromosome of clinical and environmental clones of *V. cholerae* was surprising since the XerD-arm of TLCΦ *attP* contains 8 bp difference from *dif1*, which almost completely abolishes XerD binding (10, 11).

TLCΦ relies on its own XerD activation factor for integration, XafT (10, 23). No other *V. cholerae* or TLCΦ protein or sequence factors are required to promote a complete Xer recombination reaction between TLCΦ *attP* and *dif1 in vitro* (23). Those results suggested that XafT might itself open up the possibility to recombine TLCΦ *attP* and *dif1*.

### XafT promotes the formation of and stabilizes synaptic complexes

Our analysis of the influence of the XerD-arm of TLCΦ *attP* on the integration efficiency and stability of mini-TLCΦ plasmids showed that FtsK only promotes the recombination of *dif1* with a very small subset of the different 190 528 TLCΦ *attP* derived sites we studied, those that were the most similar to *dif1* (Figure 3, S3, 4 and S4). In contrast, XafT promoted the recombination of *dif1* with any of the 190 528 *attP* sites that carried the same residues as *dif1* at the 5^th^ position of the XerD-arm (Figure 3 and S3). In addition, it alleviated differences in the integration frequency of *attP* sites harbouring more or less canonical XerD-arms (Figure 3 and S3). Our results further indicated that FtsK did not perturb XafT-driven integration and excision events (Figure 3, S3 and 4). Finally, *in vitro* comparison of the efficiency with which XafT and FtsKγ drove recombination reactions between two *dif1* sites or between *dif1* and TLCΦ *attP* showed that TLCΦ attP/ *dif1* synaptic complexes are normally too rare and/or too transient to allow for FtsKγ-driven activation of XerD (Figure 5). Taken together, those results suggest that XafT can promote the efficient recombination of *dif1* with TLCΦ *attP-*derived sites harbouring a non-canonical XerD-arm because it helps assemble and stabilize synaptic complexes. As XafT contains a dimerization domain and directly interacts with XerD, it is tempting to propose that it recruits a XerD recombinase in *trans* at the *dif1* locus by forming a proteinaceous bridge with the XerD recombinase bound in *cis* (23). The resulting *dif1*/XerC/XerD/XafT/XerD complex could then be engaged in a larger nucleoprotein with an *attP* site solely bound by XerC via the interactions of recombinases. Thus, a limited amount of homology between the XerD-arm of the *attP* site and the canonical XerD binding site would be required to ensure the formation of recombination synapse.

### Role of the G base at 5^th^ innermost position of the XerD and XerC arms

Our NGS data allowed us to revisit the importance of the nature of the different positions of the XerD arm of *dif* sites. Whether integration was driven by FtsK or by XafT, the mini-TLCΦ plasmids that integrated the most efficiently were those that harboured the most *dif-like* motifs (Figure 3 and S3). However, the 5 outermost positions of the XerD-arm were less constrained than the 6 innermost positions (Figure 3 and S3). In addition, the possibility for XafT-driven integration was mainly influenced by the presence of a guanine at the 5^th^ position of the top strand of the XerD-arm of the *attP* sites, with XafT promoting the integration of most if not all of the plasmids that carried this base (Figure 3 and S3). The nature of the 5^th^ position of the XerD-arm of the *attP* sites was also the most important for FtsK-driven integration (Figure 3 and S3). Correspondingly, it was previously observed in *E. coli* that a *dif* site carrying a single mutation at the 5^th^ position of the XerD-arm was less proficient than wildtype *dif* for Xer recombination (35). In addition, methylation interference analysis showed that *E. coli* XerD interacted with the guanine at the 5^th^ position of the top strand of its binding site (35).

It was recently reported that *Bacillus subtilis* and *Staphylococcus aureus* XerD unload bacterial SMC complexes (36). This activity is independent from XerC. It relies on the binding of XerD to additional chromosomal loci other than *dif*. Those loci harbour a *dif-like* site composed of a bona fide XerD binding site, a 5-6bp degenerate central region and the 5 innermost bp of the XerD binding site (36). Our results raise the possibility that those sites evolved to permit the cooperative binding of two XerD molecules.

The specificity of binding of XerC and XerD is ensured by the bp composition of the 6^th^, 7^th^ and 9^th^ to 11^th^ non-palindromic positions of their binding arms (37, 38). However, contacts between *E. coli* XerC and *dif* were shown to be mainly limited to the 7 innermost bp of its binding site (35). Correspondingly, only the innermost region of the XerC-arm of *dif* sites is conserved (Figure S1B). The 5 innermost positions of the XerC- and XerD-arms are identical (Figure 1B and S1B). As XerC and XerD are highly related (Figure S1A), it is reasonable to argue that the guanine at the 5^th^ position plays the same primary role for binding of XerC as it does for binding of XerD (Figure 3 and S3). This observation could explain the apparent inefficiency of integration of CTXΦ in non-toxigenic environmental *V. cholerae* strains since both *difA* and the CTXΦ *attP* contain a non-canonical T/A bp at the 5^th^ position of their XerC-arm (Figure S1B, (11)). By extrapolation, we propose that XafT can promote the integration of TLCΦ into *difA* because it can stabilize the formation of recombination synapses between non-canonical sites (10).

Alternative Xer machineries composed of a single Xer recombinase are found in a few bacterial and archaeal species (39–41). The recombinases of those machineries are relatively distant from XerC and XerD (Figure S1A) and target sites that significantly deviate from classical *dif* sites (Figure S1B). A GC bp is also present at the 5th innermost position of the binding sites of alternative Xer recombinases (Figure S1B). It was shown to be important for the binding and activity of *H. pylori* XerH (39). However, this bp is inverted with respect to the GC bp of canonical XerC and XerD-arms (Figure S1B). X-ray structure analysis revealed that *H. pylori* XerH contacts the guanine of the bottom strand of its binding site with an arginine of its N-terminal domain, R65 (39). In contrast, a model based on the crystal structure of *E. coli* XerD and the Catabolite Activator Protein-DNA complex suggested that XerD contacts the guanine of the top strand of its binding site with a conserved arginine of its C-terminal domain (R220 in *E. coli* XerD and R224 in *V. cholerae* XerD, (42)). Those observations highlight the evolutionary distance between conventional and alternative Xer machineries.

### XerD-arm degeneracy prevents FtsK-driven Xer-mediated excision events

The analysis of the stability of the XafT^+^ n_8_gtg, n_5_a_2_n_2_g and tagn_8_ plasmids that were integrated at the *dif1* locus, which corresponds to the normal location of TLCΦ in the genome of *V. cholerae* pathogenic clones of the current pandemic, revealed that the mini-TLCΦ plasmids that were efficiently integrated by FtsK were also the less stable (Figure 4 and S4). In addition, all of the 190 528 mini-TLCΦ plasmids we studied, including those that harboured the *attP* sites that most resembled *dif1* were far more stable that a DNA cassette flanked by two *dif1* sites (Figure 4 and S4). Those observations lend support to the idea that the few non-canonical bp in the XerD-arm of the GGI and TLCΦ *attP* sites serve to prevent their excision (24). Nevertheless, TLCΦ can excise from its host genome (10). Our results suggest that those excision events are promoted by XafT independently of FtsK (Figure 4). They are extremely rare in normal laboratory growth conditions, but they can be increased in conditions that mimic cholera, which suggest that the production of XafT is tightly regulated (Figure 4).

### Altering the possibility for synapse formation is a major source of control

The dimer resolution sites of multicopy plasmids exploiting Xer and the attachment sites of IMEX relying on the non-canonical XerC-first recombination pathway contain a few degenerate positions in the XerC and XerD arms, which could limit the efficiency of recombination (Figure S1B). Our results suggest that those mutations evolved to limit the formation of and/or stability of synaptic complexes, thereby preventing the formation of multimers by Xer recombination (43). In particular, there are two mutations in the inner region of the XerC-arm of the ColE1 dimer resolution site, *cer*, including one at the 5^th^ position, and there is a mutation at the 6^th^ position of its XerD-arm (Figure S1B). The efficiency of intramolecular dimer resolution events relies on accessory proteins that bind to accessory sequences flanking the core *cer* site, which bring together the core *cer* sites in a synaptic complex in a specific topological configuration (22, 44). It remains to be determined whether IMEX such as CTXΦ or VGJΦ simply rely on the amplification of their free form by replication to achieve efficient integration or whether unknown mechanisms favour the formation of recombination synapses between the host target *dif* site and their *attP* sites (45, 46). Future studies will also need to address the evolutionary pressures that seems to have set the non-canonical bp of each plasmid dimer resolution and IMEX attachment sites.

## ACKNOWLEDGEMENTS

We thank James Provan, Raphaël Guérois and Virginia Lioy for helpful discussions. The work was supported by the ERC (FP7/2007-2013, grant number 28159), the ANR (grants 2016-CE12-0030-0 and 2018-CE12-0012-03) and the FRM (EQU202003010328).

## MATERIALS AND METHODS

### Strains, plasmids and oligonucleotides

Strains, plasmids and oligonucleotides are listed in Supplementary Tables S2, S3 and S4, respectively. Strains were built by natural transformation using appropriate selection markers and/or blue/white β-galactosidase screens. Plasmids pSM11 and pSM15 are pTLC8 and pSM12 derivatives, respectively (23). They were constructed by replacing the TLCΦ *attp* locus by the BsaI-*ccdB*-BsaI cassette of pFB5 (47) using Gibson assembly (48). Pools of n_8_gtg, tagn_8_ and n5a2g3 TLCΦ *attp-*derived sites were built by primer extension of oligo 4069 annealed to degenerate 4066-4068 oligonucleotides. The recombination site libraries were inserted in pSM11 and pSM15, and cloned in FCV14. Transformation reactions were plated on 20 cm diameter petri dishes and repeated to obtain a total of about 10^6^ colonies. Plasmid libraries were then extracted from ~1×10^10^ FCV14 cells, and transformed into β2163.

### Integration assays

*V. cholerae* recipients were grown to an OD600 of 0.3 in LB with 0.2% of L-arabinose (L-ara), to induce the Xer machinery. *E. coli* β2163 donors were grown to an OD600 of 0.6 in LB supplemented with 0.3 mM of Dap, and mixed with recipients at a 1:10 ratio. Donor and recipient cells were incubated for 3h on LB agar plates supplemented with Dap and L-ara, resuspended in LB supplemented with 0.2% of L-ara and incubated for an additional 1h. Conjugants were selected for the plasmid antibiotic resistance and Dap autotrophy. In the case of the degenerate plasmid libraries, several conjugations were performed for each integration assay to ensure the recovery of ~10^6^ clones.

### Stability assays

To observe the evolution of the different *attP* classes during the proliferation FtsK and XafT, integration libraries were incubated without any selection pressure for the maintenance of the integrated elements. Fresh LB was inoculated with ~1×10^9^ cells from the integration libraries, with the addition of arabinose 0.2% for the expression of XerD and XerC.

### NGS analysis

Plasmid *attP* sites were amplified by performing 17 PCR cycles on 50-100 ng of plasmid library gDNA with an equimolar mix of 4103-4105 P5 and 4178-4180/4246-4248 P7 primers. The products were purified from the P5 and P7 primers the double selection with AMPure. NextSeq reads were trimmed with Cutadapt (version 1.17). For the 2D-maps, different [x y] coordinates were assigned to each nucleotide for the 65 536 possible motifs as follows: the [x, y] coordinates were initially set to [0 0]. Then, [1/2n-1, 1/2 n-1], [−1/2n-1 /2n-1], [−1/2n-1, −1/2n-1] or [1/2n-1, −1/2n-1] were added to [x, y] for each n position of the degenerate motif if the base of the recombination site was A, T, G or C, respectively.

### *In vitro* recombination assays

Proteins were purified and as *In vitro* recombination assays were performed as described in (23).

## SUPPLEMENTARY FIGURE LEGENDS

**Supplementary Figure S1.**
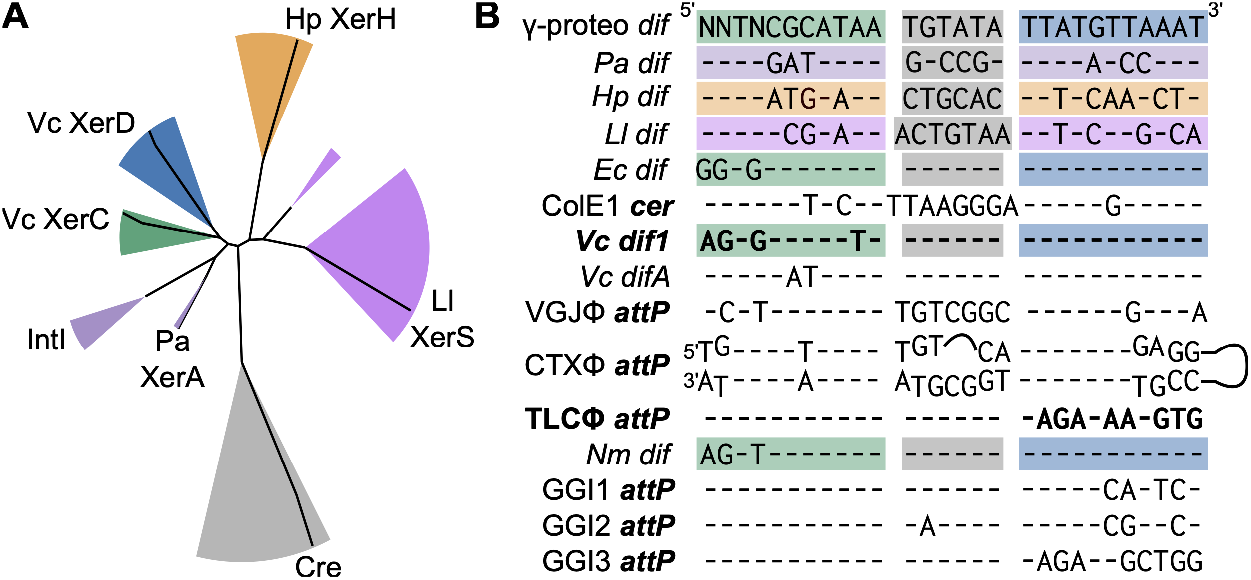
Xer recombination. (A) Phylogeny of Xer recombinases. Grey, Green, Blue, Orange, Pink and Magenta sectors: Cre (outgroup), XerC, XerD, XerH, XerS, XerA and IntI families, respectively. Vc: *V. cholerae;* Hp: *Helicobacter pylori;* Ll: *Lactoccocus lactis;* Pa: *Pyrococcus abessi*. (B) Sequence of the top strand of bacterial *dif* sites and of Xer recombination sites harboured by typical mobile genetic elements. The top strand is the site cleaved by XerC during recombination. Grey: central region; Green, Blue, Magenta, Orange and Pink: XerC-, XerD-, Pa XerA-, Hp XerH- and Ll XerS-binding arms, respectively. γ-proteo: γ-proteobacteria *dif* consensus. ColE1 *cer*: core of the ColE1 plasmid dimer resolution site. Attachment site bases homologous to the host *dif* sequence are indicated by a hyphen. CTXΦ *attP* is the stem of a folded hairpin, with 12 nt on the top strand of its central region and 7 on its bottom strand. Nm: *Neisseria meningitidis*.

**Supplementary Figure S2.**
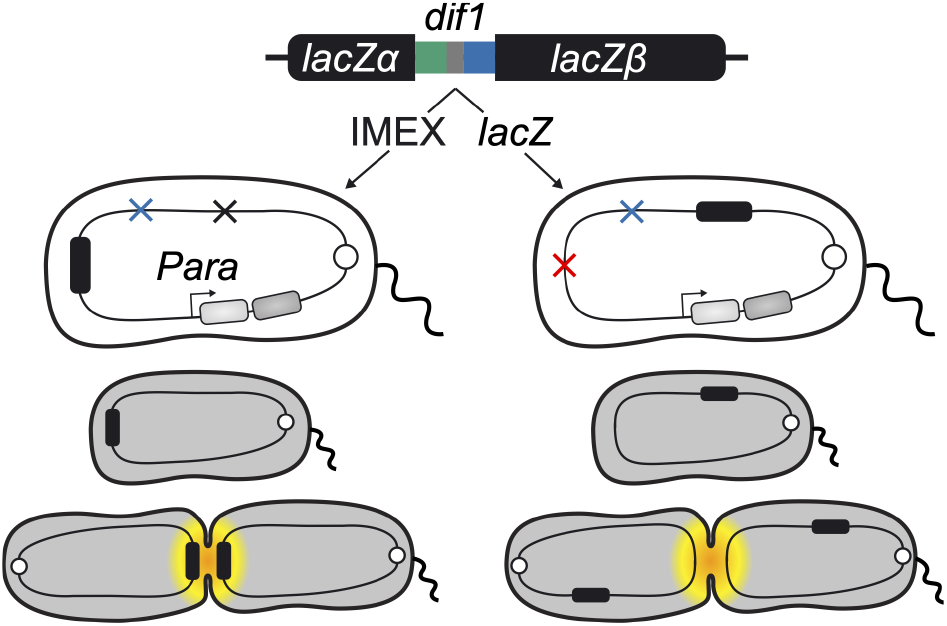
Parallel monitoring of integration and stability. Scheme of the V. cholerae reporter strains. White circle: chromosome 1 replication origin; red, blue and black crosses: IMEX, xerD and lacZ deletions, respectively; black rectangle: E. coli lacZα-dif1-lacZβ gene; light and dark grey rectangles: synthetic xerC and xerD operon under the control of the arabinose promoter (Para). Grey and yellow shadings depict the subcellular region and timing of activity of FtsK during the cell cycle.

**Supplementary Figure S3.**
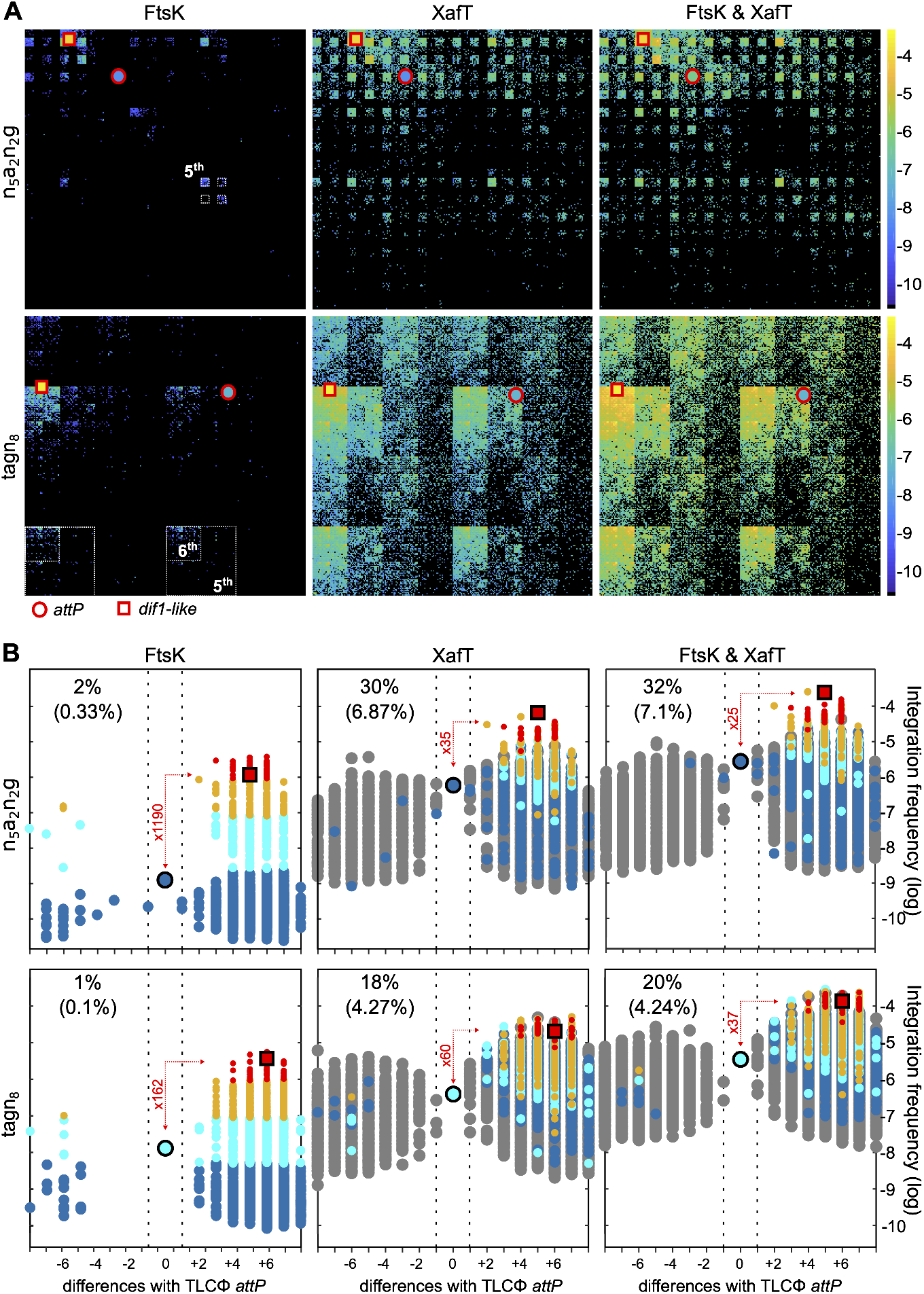
Suicide mini-TLCΦ plasmid integration. **(A)** 2D maps representation of the relative integration efficiency of the different possible sequences. **(B)** Butterfly plot representation of the relative integration efficiency of the different possible sequences. Legend as in Figure 3.

**Supplementary Figure S4.**
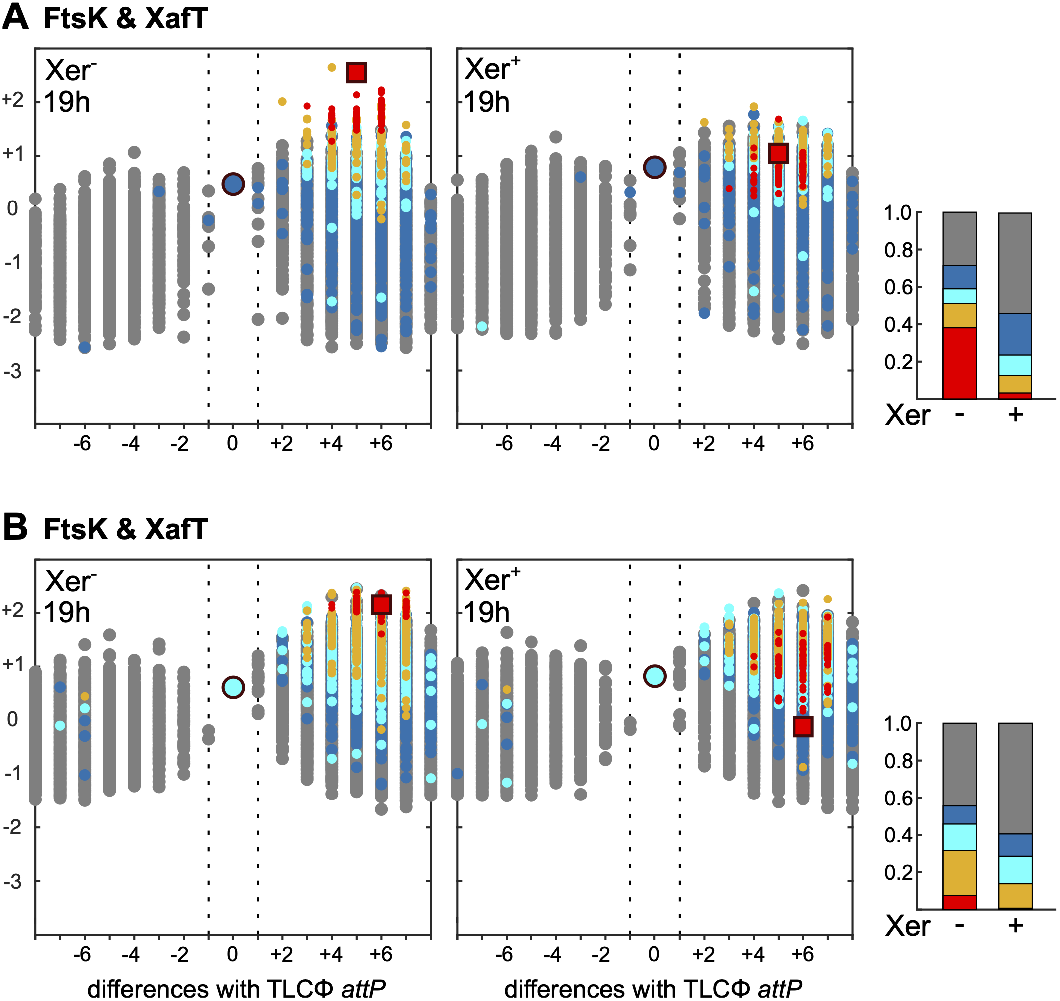
Relative stability of XafT^+^ plasmids. **(A)** Remaining n_5_a_2_n_3_g plasmids at the natural *dif1* locus after growth in LB. **(B)** Remaining tagn_8_ plasmids at the natural *dif1* locus after growth in LB. (tagn_8_ motif. Legend as in Figure 4.

## SUPPLEMENTARY TABLES

**Table S1.** NGS data.

| Degenerate pool | XeD activation factor | Sequence library | Number of reads | Expected copy number | Absent motifs | % present motifs | Combined pool |  |
| --- | --- | --- | --- | --- | --- | --- | --- | --- |
|  |  |  |  |  |  |  | Absent motifs | % present motifs |
| n8gtg | XafT | Plasmid | 3014513 | 46 | 87 | 99,9% | 270 | 99,9% |
| tagn8 | XafT | Plasmid | 1375803 | 21 | 146 | 99,8% |  |  |
| n5a2n3g | XafT | Plasmid | 1003332 | 15 | 165 | 99,7% |  |  |
| n8gtg | - | Plasmid | 1500796 | 23 | 318 | 99,5% | 425 | 99,8% |
| tagn8 | - | Plasmid | 1443196 | 22 | 62 | 99,9% |  |  |
| n5a2n3g | - | Plasmid | 1433248 | 22 | 126 | 99,8% |  |  |
| n8gtg | FtsK | Integration | 865170 | 13 | 64116 | 2,2% | 185793 | 2,5% |
| tagn8 | FtsK | Integration | 784541 | 12 | 63381 | 3,3% |  |  |
| n5a2n3g | FtsK | Integration | 981722 | 15 | 64324 | 1,8% |  |  |
| n8gtg | XafT | Integration | 2083611 | 32 | 55038 | 16,0% | 142984 | 25,0% |
| tagn8 | XafT | Integration | 1684057 | 26 | 35252 | 46,2% |  |  |
| n5a2n3g | XafT | Integration | 891750 | 14 | 57896 | 11,7% |  |  |
| n8gtg | FtsK & XafT | Integration | 1469756 | 22 | 53936 | 17,7% | 137400 | 27,9% |
| tagn8 | FtsK & XafT | Integration | 1166802 | 18 | 33214 | 49,3% |  |  |
| n5a2n3g | FtsK & XafT | Integration | 1022000 | 16 | 55223 | 15,7% |  |  |
| n8gtg | FtsK & XafT | Excision Xer- 19h | 1188318 | 18 | 53784 | 17,9% | 137606 | 27,8% |
| tagn8 | FtsK & XafT | Excision Xer- 19h | 1249627 | 19 | 33001 | 49,6% |  |  |
| n5a2n3g | FtsK & XafT | Excision Xer- 19h | 1202064 | 18 | 55845 | 14,8% |  |  |
| n8gtg | FtsK & XafT | Excision Xer+ 19h | 1575885 | 24 | 52833 | 19,4% | 135158 | 29,1% |
| tagn8 | FtsK & XafT | Excision Xer+ 19h | 1274840 | 19 | 32094 | 51,0% |  |  |
| n5a2n3g | FtsK & XafT | Excision Xer+ 19h | 771612 | 12 | 55160 | 15,8% |  |  |
| n8gtg | XafT | Excision Xer- 42h | 1542591 | 24 | 55548 | 15,2% | - | - |
| n8gtg | XafT | Excision Xer+ 42h | 473362 | 7 | 56523 | 13,8% | - | - |
| n8gtg | XafT | Excision Xer- 21h | 1082288 | 17 | 55903 | 14,7% | - | - |
| n8gtg | XafT | Excision Xer- 42h | 1334926 | 20 | 55811 | 14,8% | - | - |
| n8gtg | XafT | Excision Xer- 63h | 1518529 | 23 | 55639 | 15,1% | - | - |
| n8gtg | XafT | Excision Xer+ 21h | 1085199 | 17 | 55698 | 15,0% | - | - |
| n8gtg | XafT | Excision Xer+ 42h | 1075984 | 16 | 56049 | 14,5% | - | - |
| n8gtg | XafT | Excision Xer+ 63h | 1179951 | 18 | 58085 | 11,4% | - | - |

**Table S2.** Strain list.

|  | Genotype | Reference |
| --- | --- | --- |
| CMV30 | N1696 ChapR $\Delta xerD::sh ble$ (ZeoR) $xerC::paraXerCD-SpecR \Delta lacZ dif1-prophages::EclacZa-dif1-lacZb$ | (49) |
| EPV369 | N16961 ChapR $xerC::paraXerCD-SpecR \Delta lacZ::lacZ\alpha-dif1-lacZ\beta \Delta dif1-prophages \Delta xerD::sh ble$ (ZeoR) | This study |
| EPV366 | N16961 ChapR $xerC::paraXerCD-SpecR \Delta lacZ::lacZ\alpha-dif1-lacZ\beta @ lacZ \Delta dif1-prophages$ | This study |
| FCV14 | *E. coli* $xerC::KanR$ DH5α producing the R6K $\pi$ protein | Laboratory collection |
| β2163 | *E. coli* (F-) RP4-2-Tc::Mu $\Delta dapA::(erm-pir)$ [Km<sup>R</sup> Em<sup>R</sup>] | (50) |

**Table S3.** Plasmid list.

| Plasmids | Properties | Reference |
| --- | --- | --- |
| pTLC8<br>(pCM33) | pTLC derived vector lacking all TLCΦ nucleotides from 325 to 4137 | (1) |
| pTLC10<br>(pCM24) | pTLC derived vector with a stop mutation in VC1465. | Midonet et al., 2014) |
| pTLC-Cri<sup>+</sup><br>(pBS90) | R6K γ ori, CmR, oriT plasmid carrying TLCΦ | (Midonet et al., 2014) |
| pFB0005 (pET-Gate2) | Backbone for Golden gate assembly for gene targeting in *Bacillus subtilis* | (51) |
| pSM11 | Derivative of pTLC8 in which *attP<sub>TLC</sub>* was replaced by *Bsa1-ccdb-Bsa1* | This study |
| pSM12 | Derivative of pTLC10 lacking all TLCΦ nucleotides from 325 to 4137 | This study |
| pSM15 | Derivative of pSM12 in which *attP<sub>TLC</sub>* was replaced by *Bsa1-ccdb-Bsa1* | This study |
| pSM17 | Derivative of pSM11 carrying *dif1* | This study |
| pSM18 | Derivative of pSM15 carrying *dif1* | This study |
| pCM153 | pBR322 + MBP-6His-TEVsite-VC1465 under T7 promotor and lacO operator + KanR | (23) |
| pCM157 | pBR322 + MBP-6His-TEVsite-V. *cholerae* XerC under T7 promoter and lacO operator + KanR | (23) |
| pCM162 | pBR322 + MBP-6His-TEVsite-V. *cholerae* XerD under T7 promoter and lacO operator + KanR | (23) |
| pCM163 | pBR322 + MBP-6His -TEVsite-V. *cholerae* XerC catalytic mutant (KQ mutation) under T7 promoter and lacO operator + KanR | (23) |
| pCM164 | pBR322 + MBP-6His -TEVsite-V. *cholerae* XerD catalytic mutant (KQ mutation) under T7 promoter and lacO operator + KanR | (23) |

**Table S4.**
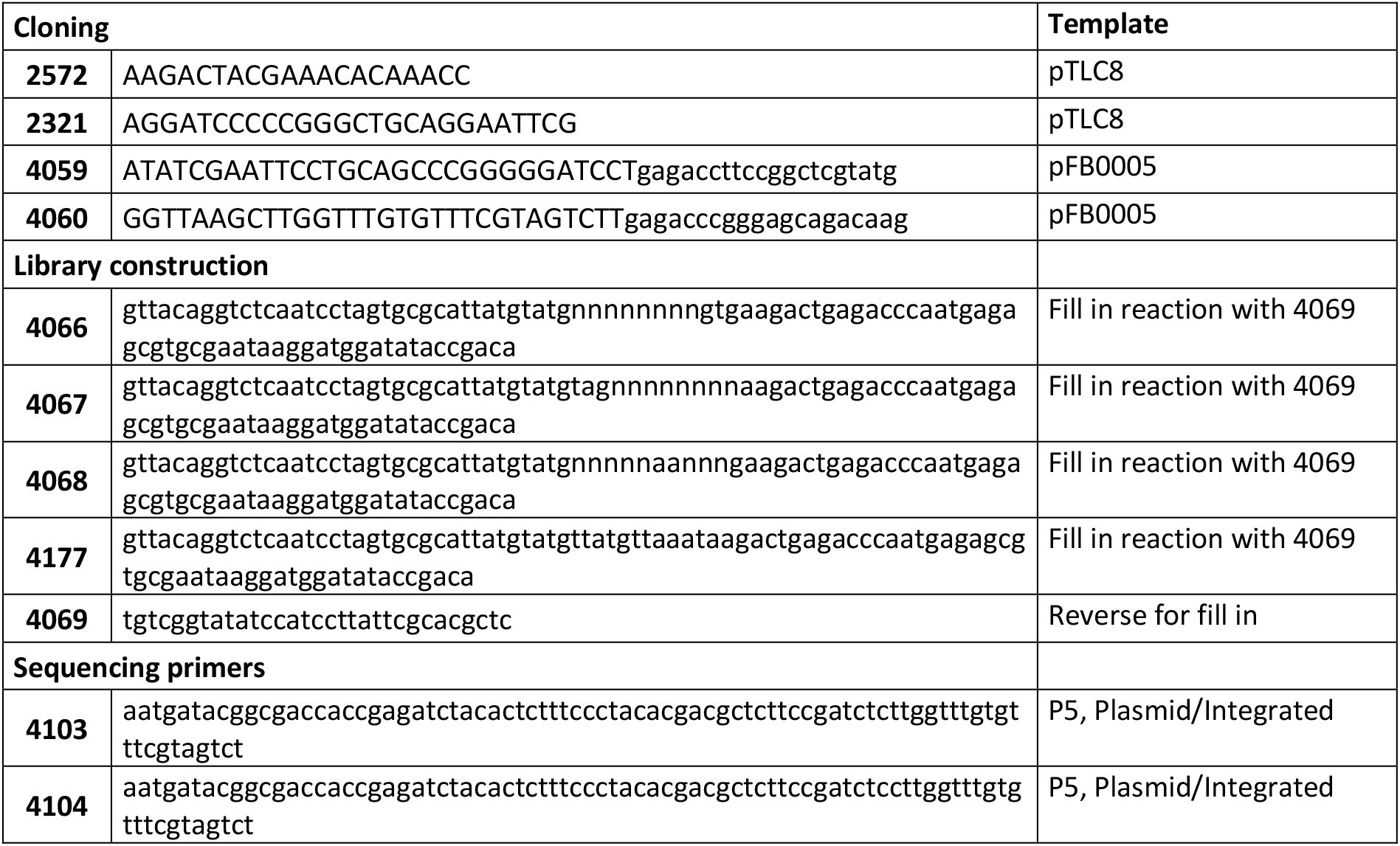
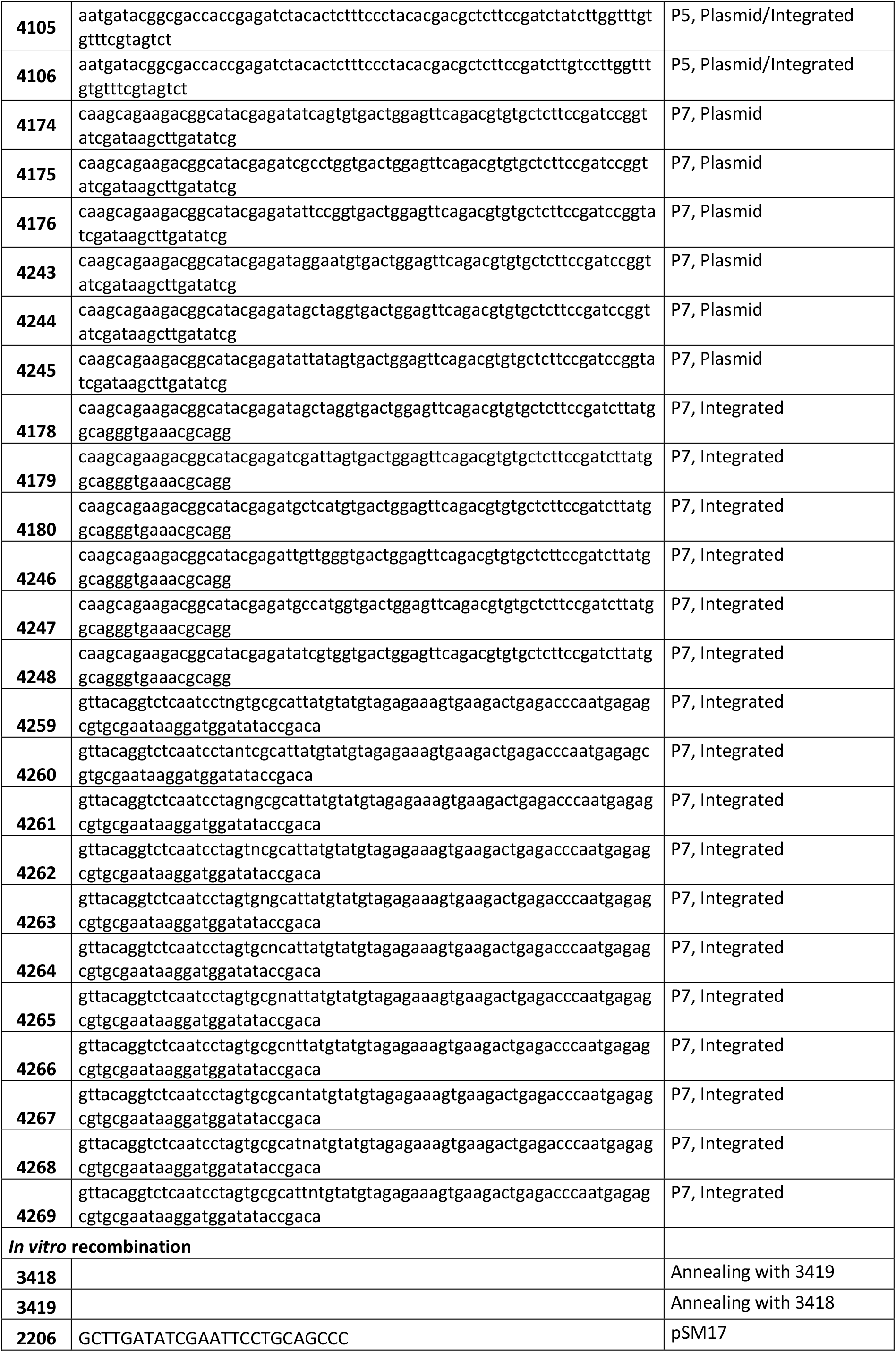
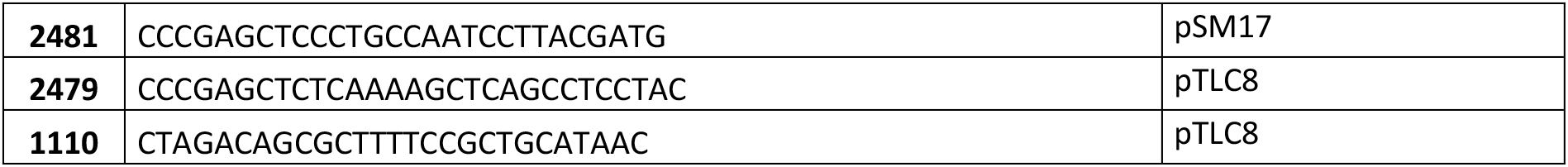
Oligonucleotide list.

## Notes

### Competing Interest Statement

The authors have declared no competing interest.

## REFERENCES

1. C. Midonet, F.-X. Barre, Xer Site-Specific Recombination: Promoting Vertical and Horizontal Transmission of Genetic Information. Microbiol. Spectr. 2 (2014).

2. G. Blakely, et al., Two related recombinases are required for site-specific recombination at dif and cer in E. coli K12. Cell 75, 351–361 (1993).

3. S. D. Colloms, P. Sykora, G. Szatmari, D. J. Sherratt, Recombination at ColE1 cer requires the Escherichia coli xerC gene product, a member of the lambda integrase family of site-specific recombinases. J Bacteriol 172, 6973–80 (1990).

4. B. Das, E. Martínez, C. Midonet, F.-X. Barre, Integrative mobile elements exploiting Xer recombination. Trends Microbiol. 21, 23–30 (2013).

5. P. Balalovski, I. Grainge, Mobilization of pdif modules in Acinetobacter: A novel mechanism for antibiotic resistance gene shuffling? Mol. Microbiol. (2020) https://doi.org/10.1111/mmi.14563.

6. M. K. Waldor, J. J. Mekalanos, Lysogenic conversion by a filamentous phage encoding cholera toxin. Science 272, 1910–1914 (1996).

7. K. E. Huber, M. K. Waldor, Filamentous phage integration requires the host recombinases XerC and XerD. Nature 417, 656–659 (2002).

8. B. Das, J. Bischerour, M.-E. Val, F.-X. Barre, Molecular keys of the tropism of integration of the cholera toxin phage. Proc. Natl. Acad. Sci. U. S. A. 107, 4377–4382 (2010).

9. B. Das, J. Bischerour, F.-X. Barre, VGJphi integration and excision mechanisms contribute to the genetic diversity of Vibrio cholerae epidemic strains. Proc. Natl. Acad. Sci. U. S. A. 108, 2516–2521 (2011).

10. C. Midonet, B. Das, E. Paly, F.-X. Barre, XerD-mediated FtsK-independent integration of TLCφ into the Vibrio cholerae genome. Proc. Natl. Acad. Sci. U. S. A. 111, 16848–16853 (2014).

11. F. Hassan, M. Kamruzzaman, J. J. Mekalanos, S. M. Faruque, Satellite phage TLCφ enables toxigenic conversion by CTX phage through dif site alteration. Nature 467, 982–985 (2010).

12. L. Aussel, et al., FtsK Is a DNA motor protein that activates chromosome dimer resolution by switching the catalytic state of the XerC and XerD recombinases. Cell 108, 195–205 (2002).

13. M.-E. Val, et al., FtsK-dependent dimer resolution on multiple chromosomes in the pathogen Vibrio cholerae. PLoS Genet. 4, e1000201 (2008).

14. J. Yates, M. Aroyo, D. J. Sherratt, F.-X. Barre, Species specificity in the activation of Xer recombination at dif by FtsK. Mol. Microbiol. 49, 241–249 (2003).

15. I. Grainge, C. Lesterlin, D. J. Sherratt, Activation of XerCD-dif recombination by the FtsK DNA translocase. Nucleic Acids Res 39, 5140–8 (2011).

16. J. Yates, et al., Dissection of a functional interaction between the DNA translocase, FtsK, and the XerD recombinase. Mol Microbiol 59, 1754–66 (2006).

17. S. Bigot, et al., KOPS: DNA motifs that control E. coli chromosome segregation by orienting the FtsK translocase. EMBO J. 24, 3770–3780 (2005).

18. S. P. Kennedy, F. Chevalier, F.-X. Barre, Delayed activation of Xer recombination at dif by FtsK during septum assembly in Escherichia coli. Mol. Microbiol. 68, 1018–1028 (2008).

19. F. X. Barre, et al., FtsK functions in the processing of a Holliday junction intermediate during bacterial chromosome segregation. Genes Dev. 14, 2976–2988 (2000).

20. F. Cornet, J. Louarn, J. Patte, J. M. Louarn, Restriction of the activity of the recombination site dif to a small zone of the Escherichia coli chromosome. Genes Dev 10, 1152–61 (1996).

21. M.-E. Val, et al., The single-stranded genome of phage CTX is the form used for integration into the genome of Vibrio cholerae. Mol. Cell 19, 559–566 (2005).

22. S. D. Colloms, J. Bath, D. J. Sherratt, Topological selectivity in Xer site-specific recombination. Cell 88, 855–864 (1997).

23. C. Midonet, S. Miele, E. Paly, R. Guerois, F.-X. Barre, The TLCΦ satellite phage harbors a Xer recombination activation factor. Proc. Natl. Acad. Sci. 116, 18391–18396 (2019).

24. F. Fournes, et al., FtsK translocation permits discrimination between an endogenous and an imported Xer/dif recombination complex. Proc. Natl. Acad. Sci. U. S. A. 113, 7882–7887 (2016).

25. E. Galli, C. Midonet, E. Paly, F.-X. Barre, Fast growth conditions uncouple the final stages of chromosome segregation and cell division in Escherichia coli. PLoS Genet. 13, e1006702 (2017).

26. E. Espinosa, E. Paly, F.-X. Barre, High-Resolution Whole-Genome Analysis of Sister-Chromatid Contacts. Mol. Cell 79, 857–869.e3 (2020).

27. G. Demarre, et al., Differential management of the replication terminus regions of the two Vibrio cholerae chromosomes during cell division. PLoS Genet. 10, e1004557 (2014).

28. A. David, et al., The two Cis-acting sites, parS1 and oriC1, contribute to the longitudinal organisation of Vibrio cholerae chromosome I. PLoS Genet. 10, e1004448 (2014).

29. J. P. Dillard, H. S. Seifert, A variable genetic island specific for Neisseria gonorrhoeae is involved in providing DNA for natural transformation and is found more often in disseminated infection isolates. Mol Microbiol 41, 263–77 (2001).

30. N. M. Domínguez, K. T. Hackett, J. P. Dillard, XerCD-Mediated Site-Specific Recombination Leads to Loss of the 57-Kilobase Gonococcal Genetic Island. J. Bacteriol. 193, 377–388 (2011).

31. M. Iwanaga, K. Yamamoto, New medium for the production of cholera toxin by Vibrio cholerae O1 biotype El Tor. J. Clin. Microbiol. 22, 405–408 (1985).

32. A. N. Keller, et al., Activation of Xer-recombination at dif: structural basis of the FtsKγ-XerD interaction. Sci. Rep. 6, 33357 (2016).

33. T. H. Massey, L. Aussel, F.-X. Barre, D. J. Sherratt, Asymmetric activation of Xer site-specific recombination by FtsK. EMBO Rep. 5, 399–404 (2004).

34. L. Bonné, S. Bigot, F. Chevalier, J.-F. Allemand, F.-X. Barre, Asymmetric DNA requirements in Xer recombination activation by FtsK. Nucleic Acids Res. 37, 2371–2380 (2009).

35. G. W. Blakely, D. J. Sherratt, Interactions of the site-specific recombinases XerC and XerD with the recombination site dif. Nucleic Acids Res. 22, 5613–5620 (1994).

36. X. Karaboja, et al., XerD unloads bacterial SMC complexes at the replication terminus. Mol. Cell 81, 756–766.e8 (2021).

37. G. Blakely, D. Sherratt, Determinants of selectivity in Xer site-specific recombination. Genes Dev. 10, 762–773 (1996).

38. F. Hayes, D. J. Sherratt, Recombinase binding specificity at the chromosome dimer resolution site dif of Escherichia coli. J Mol Biol 266, 525–37 (1997).

39. A. Bebel, E. Karaca, B. Kumar, W. M. Stark, O. Barabas, Structural snapshots of Xer recombination reveal activation by synaptic complex remodeling and DNA bending. eLife 5.

40. P. Le Bourgeois, et al., The Unconventional Xer Recombination Machinery of Streptococci/Lactococci. PLoS Genet 3, e117 (2007).

41. I. G. Duggin, N. Dubarry, S. D. Bell, Replication termination and chromosome dimer resolution in the archaeon Sulfolobus solfataricus. EMBO J. 30, 145–153 (2011).

42. H. S. Subramanya, et al., Crystal structure of the site-specific recombinase, XerD. EMBO J 16, 5178–87 (1997).

43. D. K. Summers, C. W. Beton, H. L. Withers, Multicopy plasmid instability: the dimer catastrophe hypothesis. Mol Microbiol 8, 1031–8 (1993).

44. M. Bregu, D. J. Sherratt, S. D. Colloms, Accessory factors determine the order of strand exchange in Xer recombination at psi. EMBO J. 21, 3888–3897 (2002).

45. J. Bischerour, C. Spangenberg, F.-X. Barre, Holliday junction affinity of the base excision repair factor Endo III contributes to cholera toxin phage integration. EMBO J. 31, 3757–3767 (2012).

46. E. Martínez, E. Paly, F.-X. Barre, CTXφ Replication Depends on the Histone-Like HU Protein and the UvrD Helicase. PLoS Genet. 11, e1005256 (2015).

47. C. Engler, R. Kandzia, S. Marillonnet, A one pot, one step, precision cloning method with high throughput capability. PloS One 3, e3647 (2008).

48. D. G. Gibson, et al., Enzymatic assembly of DNA molecules up to several hundred kilobases. Nat. Methods 6, 343–345 (2009).

49. E. Galli, et al., Cell division licensing in the multi-chromosomal Vibrio cholerae bacterium. Nat. Microbiol. 1, 16094 (2016).

50. G. Demarre, et al., A new family of mobilizable suicide plasmids based on broad host range R388 plasmid (IncW) and RP4 plasmid (IncPα) conjugative machineries and their cognate Escherichia coli host strains. Res. Microbiol. 156, 245–255 (2005).

51. M.-L. Diebold-Durand, F. Bürmann, S. Gruber, “High-Throughput Allelic Replacement Screening in Bacillus subtilis” in SMC Complexes: Methods and Protocols, Methods in Molecular Biology., A. Badrinarayanan, Ed. (Springer, 2019), pp. 49–61.

